# Small molecule screen employing patient-derived iPS hepatocytes identifies LRRK2 as a novel therapeutic target for Alpha1 Antitrypsin Deficiency

**DOI:** 10.1101/2021.09.17.460732

**Authors:** Deniz Kent, Soon Seng Ng, Payam Khoshkenar, Adam M. Syanda, Chao Zheng Li, Marina Zieger, Cindy Greer, Stephanie Hatch, Joe Segal, Samuel J.I. Blackford, Vivek Chowdary, Taylor Ismali, Davide Danovi, Sunil Sahdeo, Daniel Ebner, Christian Mueller, S. Tamir Rashid

## Abstract

Alpha-1 antitrypsin deficiency is a life-threatening condition caused by inheritance of the *SERPINA1* gene ‘Z’ variant. This single base pair mutation leads to protein misfolding, ER entrapment and gain of toxic function. Despite the significant unmet medical need presented by this disorder, there remain no approved medicines and the only curative option is liver transplantation. We hypothesized that an unbiased screen of human hepatocytes harbouring the Z mutation (ATZ) using small molecules targeted against protein degradation pathways would uncover novel biological insights of therapeutic relevance. Here we report the results of that screen performed in a patient-derived iPSC model of ATZ. Starting from 1,041 compounds we identified 14 targets capable of reducing polymer burden, including Leucine-rich repeat kinase-2 (LRRK2), a well-studied target in Parkinson’s. Genetic deletion of LRRK2 in ATZ mice reduced polymers and associated fibrotic liver disease leading us to test a library of commercially available LRRK2 kinase inhibitors in both patient iPSC and CHO cell models. One of the molecules tested, CZC-25146, reduced polymer load, increased normal AAT secretion and reduced inflammatory cytokines with pharmacokinetic properties supporting its potential use for treating liver diseases. We therefore tested CZC-25146 in the ATZ mouse model and confirmed its efficacy for polymer reduction without signs of toxicity. Mechanistically, in both human and mouse models, our data show CZC-25146 inhibits LRRK2 kinase activity and induces autophagy. Cumulatively, these findings support the use of CZC-25146 and LRRK2 inhibitors in general in hepatic proteopathy disease research and as potential new treatment approaches for patients.

**One Sentence Summary:** A small molecule screen in patient iPSCs with *in vivo* validation in mice identifies LRRK2 as a new therapeutic target for Alpha-1 Antitrypsin Deficiency.

## INTRODUCTION

The ability of a cell to tightly regulate protein quality is fundamental to life *(1)*. Failure of these processes can result in accumulation of protein aggregates (proteopathy) with devastating repercussions *(2)*. Accumulation of α-1-antitrypsin (AAT) protein aggregates is an archetypal example of how dysregulation in these mechanisms can lead to life threatening human disease. In healthy individuals, AAT is a 52 kDa glycoprotein product of the *SERPINA1* gene secreted from the liver’s hepatocytes to the lungs, where it acts as a neutrophil elastase inhibitor. The ‘Z’ variant describes a single base pair mutation in Exon 5 of the *SERPINA1* gene, that results in an amino acid substitution (Glutamate for Valine) at position 342 *(3)*. This substitution leads to structural changes in the hinge region of the AAT protein, altered folding dynamics and predisposition to aggregation of retained polymeric protein (termed ATZ) within the endoplasmic reticulum. Toxic accumulation of intra-hepatocytic ATZ in turn triggers inflammation, fibrosis and downstream liver cancer and/or liver failure. Lack of AAT reaching the lung leads to uninhibited neutrophil elastase mediated parenchymal destruction and emphysema. Epidemiological studies suggest that there may be over 100,000 homozygous (ZZ) and over 100 million heterozygous (MZ, SZ) patients suffering with associated liver disease *(4)*. Despite significant advances in our understanding of why ATZ accumulation causes pathology, there remains no approved medicine to address this problem, with liver transplantation still the only curative treatment option. This represents a significant unmet medical need and requires urgent attention.

To address this challenge, we rationalized that whilst the clinical manifestations of proteopathies are often distinct from one another, the mechanisms underpinning them are largely conserved *(5–7)* and that the most efficient approach to identifying new medicines in ATZ would be to combine our prior expertise in chemical screening campaigns with a human specific proteopathy disease model. We accordingly used our patient derived induced pluripotent stem cell (hiPSC) Hepatocyte *in vitro* model of ATZ *(8–12)* to screen a small molecule library of over 1,000 known annotated small molecules (provided by Janssen Research and Development in participation with the Phenomics Discovery Initiative) with the objective of identifying protein polymer lowering molecular targets. Hits from the initial screen were subsequently validated in additional patient-iPS and cell models before one candidate was taken forwards for further mechanistic and efficacy testing *in vivo* as described in the results sections below.

## RESULTS

### A high-throughput annotated screen identifies seven new targets for A1ATD

The goal of this study was to identify new therapeutics for patients with A1ATD. To this end, we hypothesized that compounds known to target protein degradation and ER stress would be efficacious in reducing alpha-1 polymer (ATZ) load *(13–15)*. Accordingly, an annotated compound library containing molecules specific to these pathways was screened against our previously developed *in vitro* iPSC disease model derived from a patient who suffered from A1ATD related liver disease, as a result of carrying the ZZ genetic mutation (PiZ, Fig. 1A) *(8, 11)*. The iPSC-hepatocyte model was miniaturized into a 384-well plate format with high ATZ and albumin expression representing the disease and hepatic phenotypes respectively (Fig. S1A). The screen was designed to identify compounds capable of reducing polymeric ATZ, whilst preserving cell viability and phenotype. We tested each compound at 10μM concentration in duplicate for 24 hours. Efficacy was interpreted as reduction in immuno-fluorescence intensity following staining with the ATZ polymer-specific monoclonal antibody (2C1) *(16)* and capture using an automated high throughput imaging and analysis pipeline (Fig. 1B). Of the 1,041 compounds screened, we identified 14 significant hits by selecting those which generated the lowest z-scores in comparison to untreated ATZ iPSC-hepatocytes (Fig. 1C and S1B). These hits all had significantly (p<0.0001) reduced polymer expression (with z-scores around −1.92 compared to untreated ATZ controls) and fell below the 95^th^ percentile of healthy controls (Fig. S1C). The combination of polymer reduction and retained cell health was used as an indication of potential therapeutic applicability. We then refined these 14 hits further by repeating the experiments in a dose-dependent manner and only including compounds that exhibited dose-dependent response, preserved viability and albumin expression and whose literature searches were consistent with biological relevance (Fig. S2). This led us to pursue the following 7 molecular targets: kinase insert domain receptor (KDR), mitogen-activated protein kinase 8 (MAPK8), cyclin-dependent kinase 2 (CDK2), fatty acid-binding protein 1 (FABP1), nuclear receptor subfamily 3 group C member 1 (NR3C1), nuclear receptor subfamily 1 group H member 2 (NR1H2) and leucine-rich repeat kinase 2 (LRRK2). To validate these targets, we purchased commercially available inhibitors (all of which were chemically different to those used in the initial screens) and tested them in a dose dependent manner in two new iPS-Hepatocyte ATZ lines (Fig. S3).

**Fig 1.**
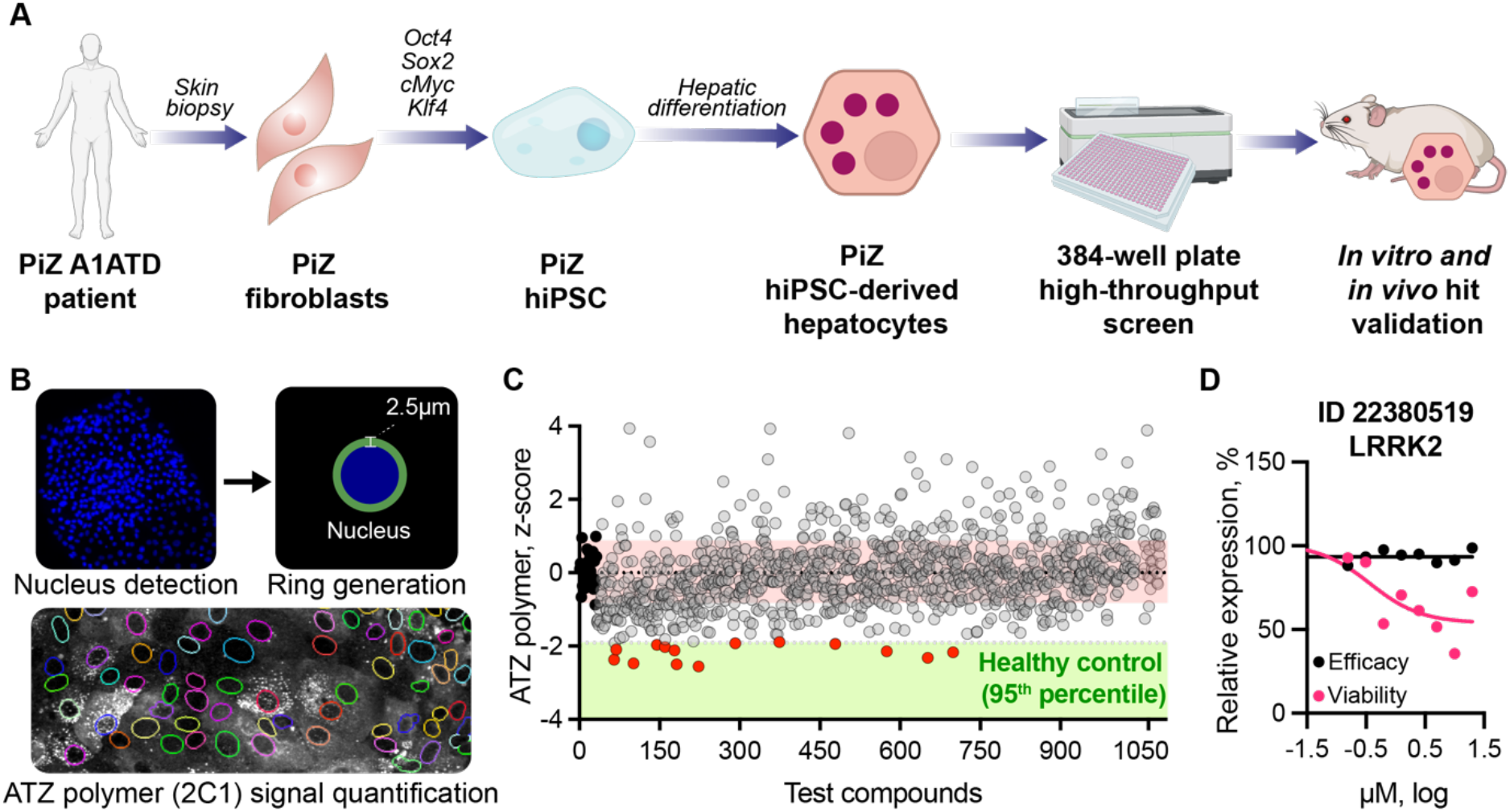
High throughput annotated compound screen in patient derived iPSC-hepatocytes identifies new targets for reducing alpha-1 antitrypsin Z (ATZ) polymers. (**A**) Schematic of the overall approach. **(B)** Image analysis pipeline involved nucleus detection and counting by DAPI signal. Each detected DAPI+ object was used to draw a 2.5um ring mask for quantification of 2C1 signal (polymeric ATZ) in each cell. (**C**) average polymer signal/cell was recorded and plotted following addition of each compound in triplicate. Reduction in polymer load (2C1) was used as a marker of therapeutic efficacy. Compounds with z-score lower than −1.92 (95^th^ percentile of healthy control cells) were selected for follow up (shown in red). Untreated A1ATD cells shown in black. Each dot represents the average of two measurements taken for each compound. N=1,041. **(D)** ID 22380519 targeting LRRK2 was identified as the top hit following the initial screen and the confirmatory dose-dependent study. It shows dose-dependent response in ATZ polymer reduction (pink) and no severe cytotoxic effects (black).

### Targeting LRRK2 reduces ATZ polymers across multiple disease models

From our initial experiments, we found inhibiting both CDK2 and LRRK2 resulted in significant polymer reduction across all patient lines (Fig. S3 and Fig. 1D). Interestingly, LRRK2 kinase inhibitors and antisense oligonucleotides are being developed as potential new medicines for another protein misfolding disease; Parkinson’s *(17)*. The combination of our experimental data and the biological relevance of this target therefore led us to further investigate the potential of LRRK2 in A1ATD as below. To begin with, we crossed the LRRK2^−/−^ mouse (a mouse whose LRRK2 gene has been mutated such that the protein product is erased) with the PiZ mouse (a mouse which expresses the Z mutant version of the human SERPINA1 gene) to generate a new mouse model (LRRK2^−/−^ x PiZ) that is both deficient in LRRK2 and pre-disposed to ATZ related liver pathology (Fig. 2A). These mice were born viable and appeared grossly healthy. Histological characterization of their livers at 8 weeks showed marked reduction in ATZ polymer load (2C1 expression) (Fig. 2, B and C) and by 28 weeks much reduced levels of fibrosis (Picrosirius red staining) (Fig. 2, D and E). These data supported our hypothesis that LRRK2 was an important regulator of ATZ polymer biology and accordingly represented an interesting therapeutic target. We therefore decided to test the polymer reducing capability of commercially available LRRK2 inhibitors, whose chemical structures targeted different binding sites of the LRRK2 kinase. For these studies, we first employed the inducible CHO cell model (a genetically modified Chinese Hamster Ovary cell which expressed the mutant Z human AAT protein following addition of Tetracycline; “Tet-ON CHO” *(18)*. Induced CHO cells were treated with one of a panel of LRRK2 inhibitors for 48 hours and their impact on polymeric ATZ levels (characterized by 2C1 immunofluorescence staining) was assessed. A dose-dependent polymer reduction (without marked toxicity) was observed with all six LRRK2 inhibitors tested except for CZC-541252 (Fig. 2F). Next, to confirm these findings in a more relevant to patient model, we repeated these studies using the optimized dose concentrations defined by the CHO experiments, in our iPSC-Hepatocyte model. Here we again observed significant polymer reduction and retention of cell health in all but one of the molecules (Fig. 2, G and H).

**Fig. 2.**
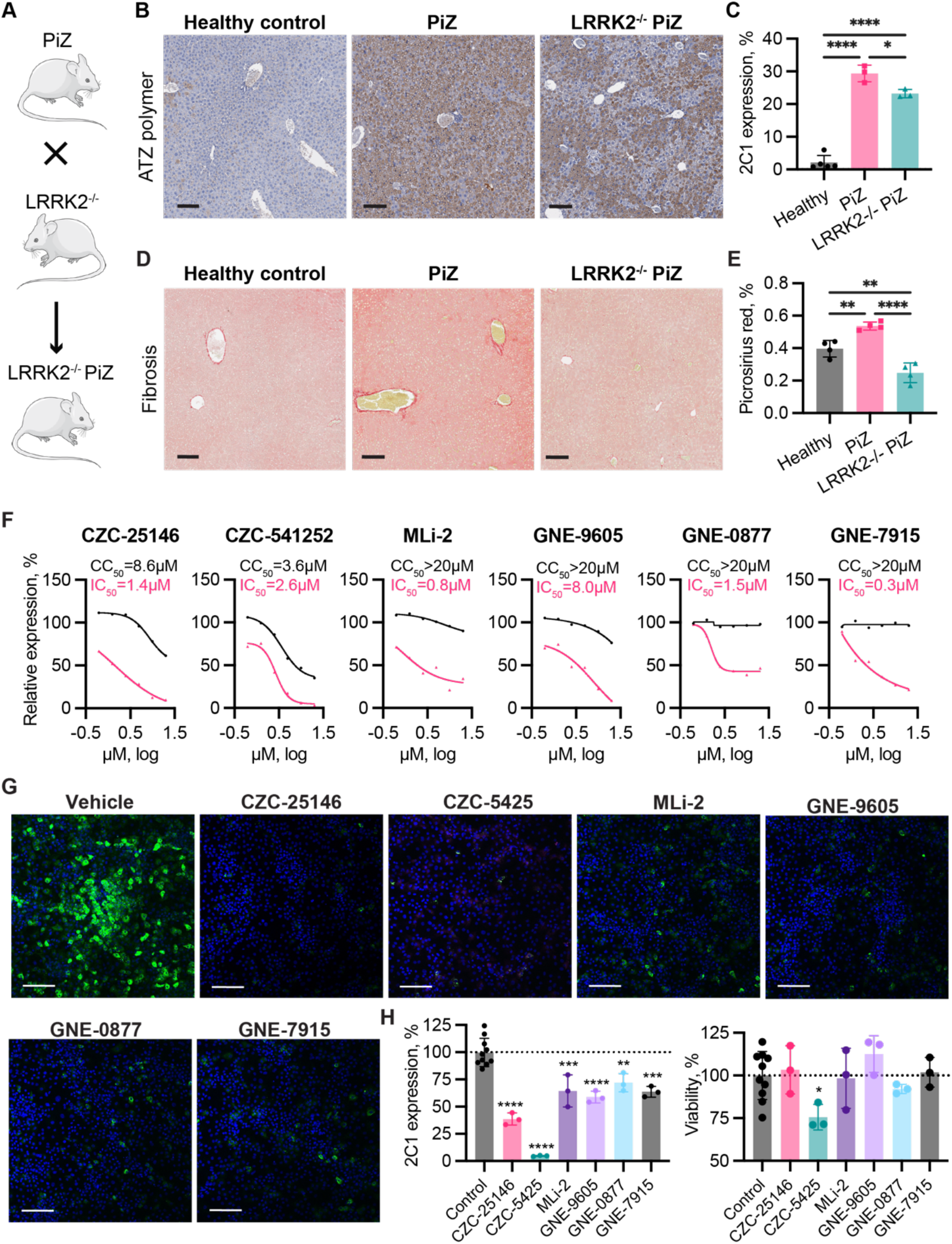
Targeting LRRK2 reduces polymeric ATZ across multiple models. (**A**) LRRK2^−/−^ PiZ mice were generated by crossing PiZ/C57B6 (expressing human ATZ polymer in liver) mice and LRRK2^−/−^/C57B6 mice. (**B**) Representative images showing ATZ polymer levels in the livers of LRRK2^−/−^ x PiZ (right), PiZ (middle) and healthy controls (left). Scale bar, 200μm. (**C**) Semi-quantitative analysis of samples from (B) shows reduced polymer in LRRK2^−/−^ x PiZ (green bars) compared to PiZ (pink bars) mice. N=3 animals. (**D**) Representative images showing fibrosis levels (Picorsirius red staining) in the livers of LRRK2^−/−^ x PiZ (right), PiZ (middle) and healthy controls (left). Scale bar, 200μm. (**E**) Semi-quantitative analysis of samples in (D) shows reduced fibrosis in LRRK2^−/−^ x PiZ (green bars) compared to PiZ (pink bars) mice. N= 4 animals. Statistical analysis by ordinary ANOVA test followed by Dunnett’s multiple comparison test. (**F**) A panel of LRRK2 inhibitors demonstrate dose-dependent polymer reduction (pink curve, inhibition) and viability (black curve, cytotoxicity) following 48 hours treatment of Tetracycline-inducible (Tet-ON) CHO-K1 cells. N=3. Nonlinear regression by four-parameter curve fit. (**G**) Representative confocal images showing polymeric ATZ expression (green) in patient iPSC-hepatocytes treated with same panel of LRRK2 inhibitors as indicated. Scale bar, 150μm. (**H**) Quantitative assessment of confocal images from (G) demonstrates significant reduction of polymer (2C1) (left) following 48h treatment without reducing cell number (viability) (right). Each dot represents a biological replicate. Statistical analysis by ordinary ANOVA test followed by Dunnett’s multiple comparison test against the control. *p<0.05, **p<0.01, ***p<0.001, ****p<0.0001.

### CZC-25146 reduces polymer load and inflammation whilst increasing secretion of AAT

Of the LRRK2 inhibitors we tested, CZC-25146 displayed the most favourable profile with regards to polymer reduction combined with cell viability *in vitro*. This compound was also noted to have poor blood-brain barrier penetrance when previously studied *in vivo (19)*. These two features led us to postulate it could be an ideal inhibitor for use in A1ATD patients with liver disease and we accordingly moved forwards with CZC-25146 to the next round of investigations.

Here we firstly observed, using three different patient derived iPSC-Hepatocyte models, that profound reductions in ATZ polymer could be achieved without compromising cell viability in a dose-dependent manner across all patient lines following 24 hours of treatment (Fig. 3, A and B and S4). After 48 hours treatment with higher doses, further reductions in ATZ polymer were seen (Fig. 3C) along with significant reductions in expression of the transcription factor nuclear factor kappa B (NFkB) (Fig. 3D), a factor previously reported to be one of the most important inducers of inflammation associated pathology in this setting *(20, 21)*. Importantly, for aiming to achieve the objective of treating both the lung (deficiency of normal AAT) and liver (toxicity of ATZ polymer) diseases known to occur with A1ATD in parallel, we observed treatment with CZC-25146 increased secretion of normal AAT at the same time as reducing intracellular ATZ polymer load (Fig. 3E). Next, to further interrogate the potential of CZC-25146 as an effective therapy for patients, we conducted a proof-of-concept study using the PiZ mouse model. The PiZ mice used in our studies were genetically modified to accumulate human Z-AAT protein in the ER of mouse hepatocytes, in a manner analogous to the pathology observed in human disease *(22)*. Notably, this animal model does not suffer from deficiency of AAT in the circulation due to endogenous production of murine AAT, thus does not recapitulate the pathology observed in the lung. Nonetheless, it is a suitable animal model to study the impact of CZC-25146 on the liver. Due to the poor water solubility of CZC-25146, absolute DMSO was used as the solvent. Equal concentrations of DMSO were administrated into animals to serve as the baseline vehicle control for analysis. Animals were dosed through oral gavage at 250mg/kg (or controls) for 14 days. We chose to use male mice exactly 6 weeks of age, which represents the peak level of polymer accumulation in this model. At 250mg/kg CZC-25146 dramatically and reproducibly reduced the polymer levels as evaluated by density of PAS-positive, diastase-resistant (PASD) globules across all five liver lobes (Fig. 4A). Inspection under higher magnification revealed marked reduction in polymeric containing globules within hepatocytes, quantitative assessment of which revealed globule reduction to be consistent across all five liver lobes (Fig. 4B top) with an overall reduction from 60% in the control group to 37% in the CZC-25146 treated group (Fig. 4B bottom). Using the Meso Scale Discovery inflammatory cytokine array, we also found that CZC-25146 administration led to a broad selection of hepatic pro-inflammatory cytokines being down-regulated and restored to wildtype levels (Fig. 4C). From a safety standpoint, no apparent drug-induced liver injury was observed following treatment, with the liver architecture remaining intact throughout lining (Fig. S5). Livers appeared to be free of fibrosis (Picrosirius Red staining) and hepatocytes did not demonstrate hyperproliferation (Ki 67) nor carcinogenesis. Lastly, even though CZC-25146 was previously defined to be a highly selective LRRK2 kinase inhibitor, activity against other kinases (PLK4, GAK, TNK1, CAMKK2, and PIP4K2C) has been postulated *in vitro (19)*. To confirm polymer reduction observed following CZC-25146 treatment of PiZ mice was a consequence of LRRK2 kinase inhibition, we harvested livers and purified protein for Western Blot (WB) analysis. LRRK2 expression and activity (as measured by autophosphorylation represented by pSer935-LRRK2), were both reduced in treated vs control mice (Fig 4D), quantification of which (Fig. 4E) was consistent with immuno-histochemical analysis (Fig. S6).

**Fig. 3.**
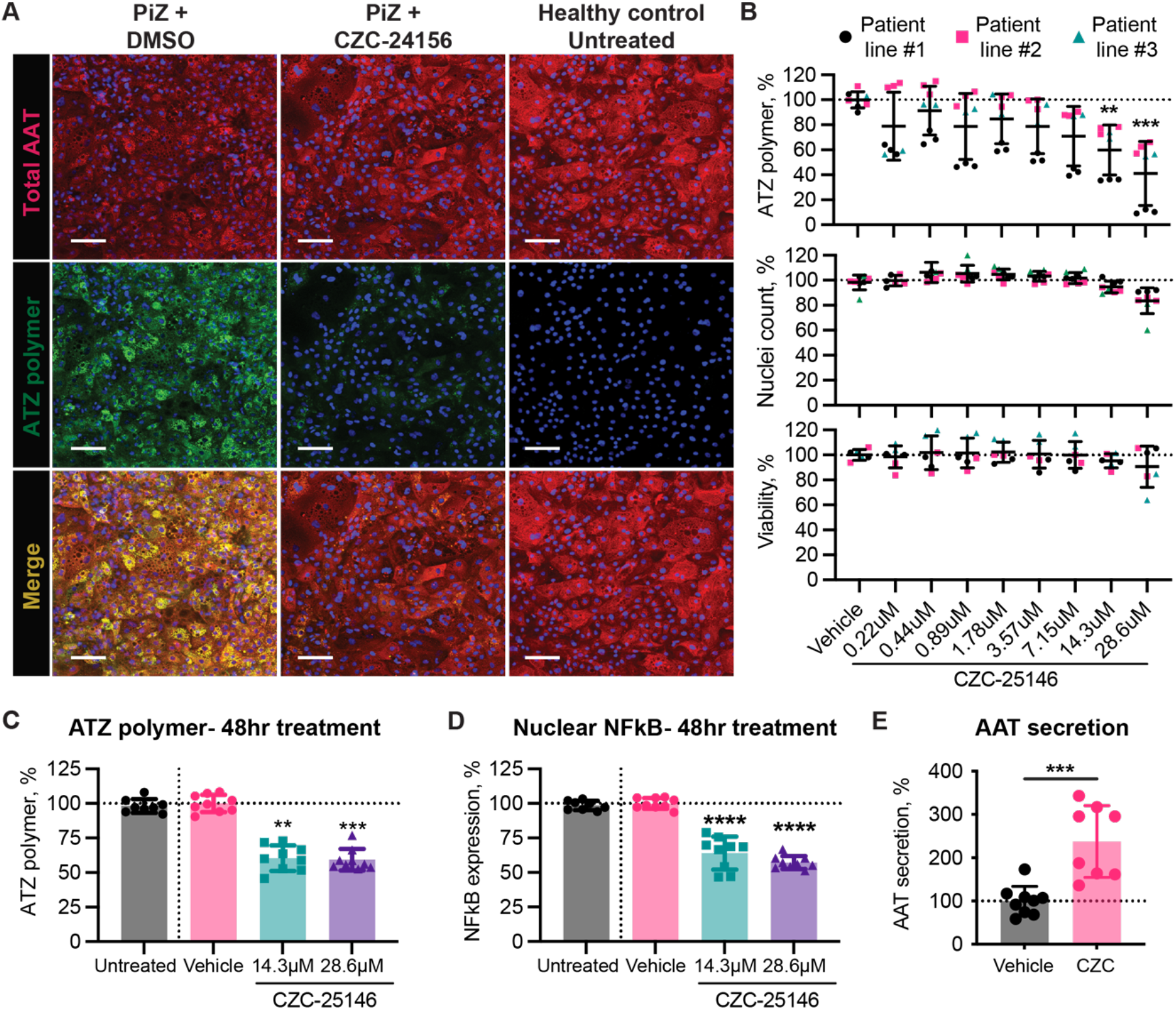
CZC-25146 reduces ATZ polymer load and restores AAT secretion *in vitro*, without compromising cell viability. (**A**) Representative images demonstrating ATZ polymer reduction in patient iPSC-Hepatocytes (middle row, green) with preserved total AAT (top row, red) after 24 hour treatment with CZC-25146 (middle column) compared to vehicle (left column) and health donor (right column) controls. Scale bar, 100μm (**B**) Quantification of polymer load (top graph), number of cells (middle graph) and cell viability (bottom graph) following increasing doses of CZC-25146 (x axis) administered for 24 hours across each of three different patient iPSC lines. 48 hour treatment of patient iPSC-Hepatocytes with CZC-25146 at high (purple bars) and lower (green bars) doses reduces intrahepatic polymeric ATZ (**C**), nuclear NFkB (**D**) and increases AAT secretion (**E**) compared to untreated (grey bars) or vehicle controls (pink bars). N=10 replicates. Age-matched healthy patient iPSC-derived hepatocytes were used as controls. Statistical analysis by Kruskal-Wallis test, followed by a Dunn’s multiple comparison test. *p<0.05, **p<0.01, ***p<0.001, ****p<0.0001.

**Fig. 4.**
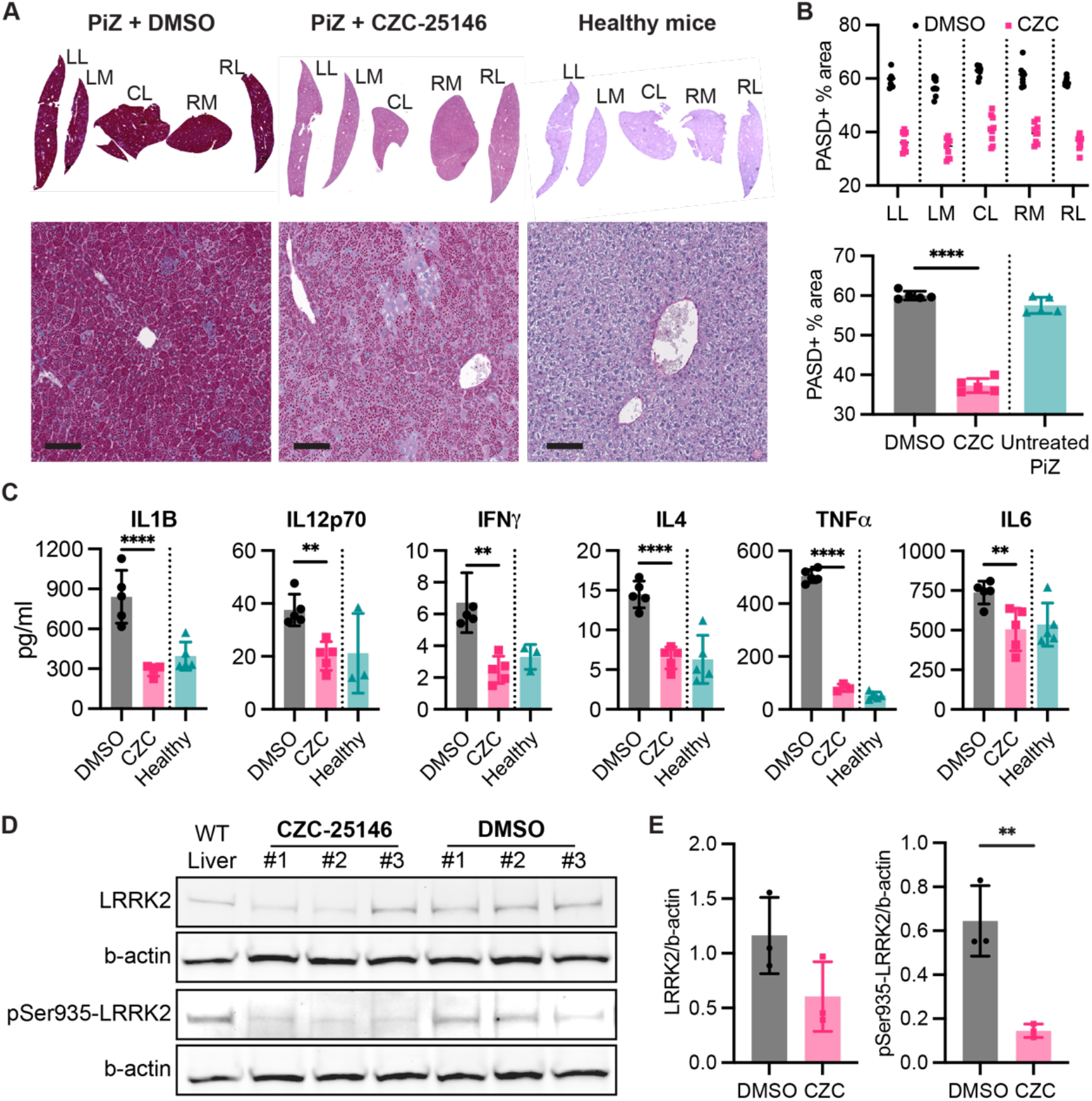
CZC-25146 reduces ATZ polymer load and decreases inflammation in the PiZ mouse liver. (**A**) Representative images of polymer load (PASD staining) in PiZ mouse livers following treatment with CZC-25146 (CZC, middle) or vehicle control DMSO (left). Healthy C57BL/6 mouse liver (right) shown as control. LL = left lateral lobe, LM = left medial lobe, CL = caudate lobe, RM = right medial lobe, RL = right lateral lobe. Scale bar, 150μm. (**B**) Quantification of staining from (A) showing reduction in each of the five liver lobules (top) and in aggregate (bottom) following treatment with CZC (pink bar), vehicle control (grey bar) or with no treatment (green bar). (**C**) levels of inflammatory cytokines in liver tissue were reduced in animals treated with CZC (pink bar) compared to vehicle control (grey bar) returning to levels seen in the no treatment (green bar) group. N=5 mice per group. (**D**) Western Blot analysis of purified LRRK2 from each of three CZC-25146 treated PiZ mice compared to vehicle only (DMSO) and untreated (WT - healthy mouse) controls showed reduction in total LRRK2 protein level (row 1) and LRRK2 phosphorylation status (row3). (**E**) Quantification analysis of blots from (D) comparing treated (pink bars) with vehicle controls (grey bars). Statistical analysis by unpaired Student t-test. N=3 mice. *p<0.05, **p<0.01, ***p<0.001, ****p<0.0001.

### CZC-25146 mediated polymer reduction is associated with autophagy induction

Dysregulated intracellular protein degradation pathways have been postulated to contribute to the liver disease associated with ATZ polymer formation *(13)*. Autophagic degradation has gained particular interest because deletion of the Atg5 gene was shown to exacerbate polymer load accumulation *(23)*. Since LRRK2 is reported to be a regulator of autophagy *(24–27)*, we hypothesized that the mechanism of effect for CZC-25146 ATZ polymer reduction could be via autophagy induction. To test our hypothesis, we first validated the functional link between CZC-25146 and autophagy, leveraging the LC3-GFP human fibroblast line, in which the autophagosome membrane is labelled with GFP. Using this assay, we found that CZC-25146 treatment induced autophagosome formation at levels similar to that observed with Rapamycin, which was used as a positive control (Fig. 5A). Time-course analysis revealed that autophagy peaked at 2 hours post-exposure to both CZC-25146 and Rapamycin, before gradually decreasing (Fig. 5B). After establishing this link between CZC-25146 and autophagy activity in fibroblasts, we next evaluated autophagy gene expression in CZC-25146 treated patient derived iPSC-Hepatocyte ATZ lines. VPS34 which is critical for the initiation of autophagy, along with downstream ATG related genes such as ATG3, ATG5, Calnexin, LC3A and LC3B were all found to be significantly upregulated following treatment, and in a dose-responsive manner (Fig. 5C). To show the functional importance of this observation, we next treated the cells with a combination of the autophagy inhibitor ammonium chloride, NH4Cl *(28)* and CZC-25146 and found that addition of NH4Cl partially reversed the polymer lowering effect of CZC-25146 (Fig. 5, D and E). Finally, to investigate whether CZC-25146 had the same effect *in vivo* we performed immuno-histochemistry for LC3 on PiZ treated mice. These analyses showed that administration of CZC-25146 could restore autophagy back towards levels seen in wild-type mice (Fig. 5, F-H).

**Fig. 5.**
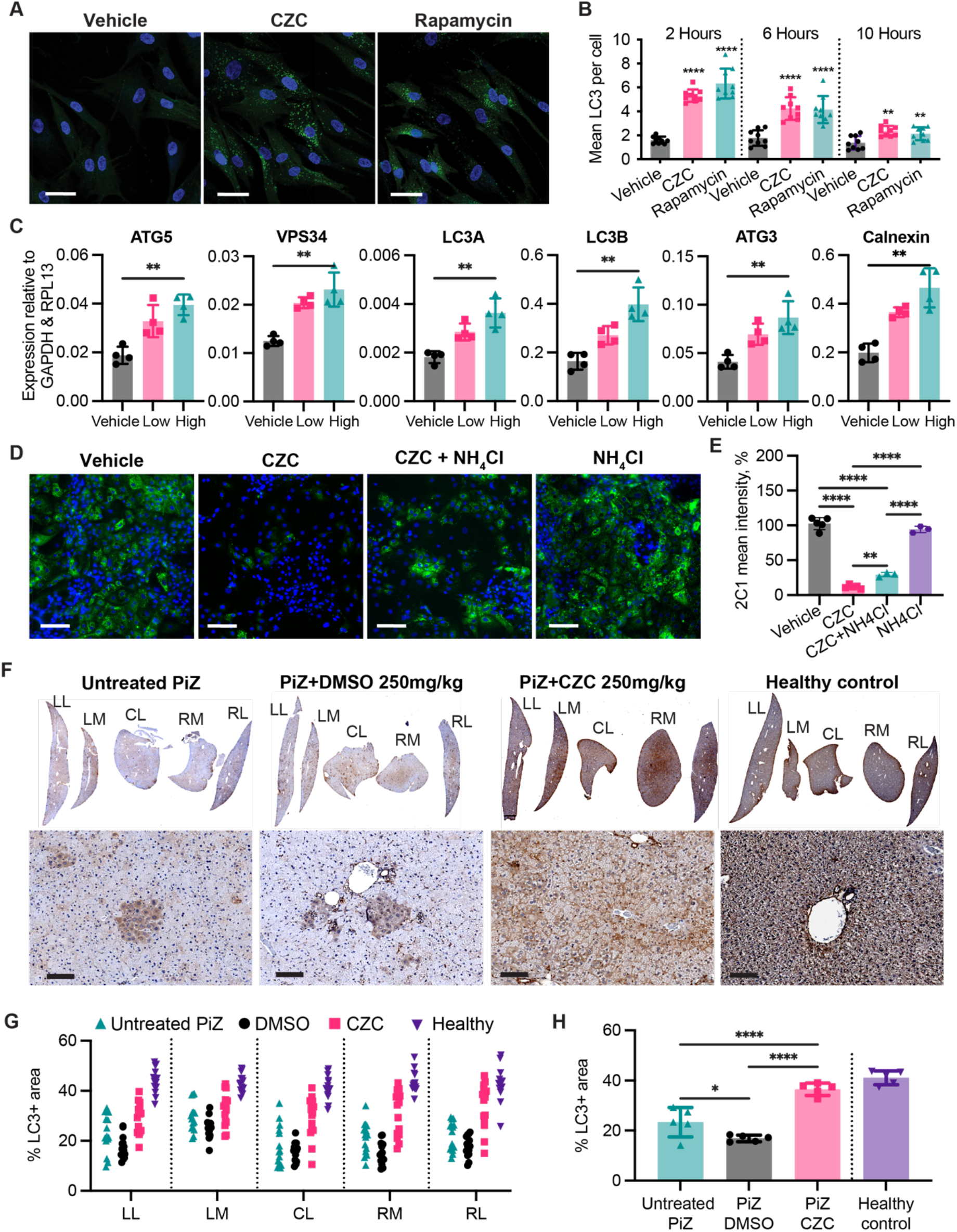
CZC-24156 induces autophagy *in vitro* and *in vivo*. (**A**) Representative images of LC3-GFP fibroblasts, following treatment with CZC-25146 (CZC, middle), vehicle control (left) or positive control Rapamcyin (right). Images taken 2 hours after exposure. Scale bar, 50μm (**B**) Time course analysis following CZC (pink) or Rapamycin (green) treatment shows significant upregulation of LC3 expression from 2-10 hours compared to vehicle controls (grey). (**C**) RT-qPCR shows dose responsive increased expression of genes involved in the autophagy pathway following addition of CZC to patient iPSC-hepatocytes at Low (pink) = 14.3μM or High (green) = 28.6μM concentrations. N=4 replicates. (**D**) Representative images showing the effect of treatment with CZC alone (left middle), CZC + autophagy inhibitor, NH4Cl (right middle), NH4Cl alone (far right) or vehicle control (far left) on ATZ polymer (2C1 expression, green) in patient iPSC-hepatocytes. Scale bar, 100μm. (**E**) Semi-quantitative analysis of images from (D). (**F**) Representative macroscopic (top) or zoomed in (bottom) images of LC3 staining in livers of PiZ (far left), healthy mice (far right), CZC-25146 treated (right middle) or vehicle control (left middle). Quantification of images from (F) separately analysed by each liver lobe (**G**) or pooled (**H**). Quantification was conducted using ImageJ. N=5 mice per group. Statistical analysis by Ordinary ANOVA test, followed by Dunnet’s multiple comparisons test. *p<0.05, **p<0.01, ***p<0.001, ****p<0.0001.

## DISCUSSION

Using a patient derived iPS Hepatocyte model of α-1-antitrypsin deficiency (A1ATD) we screened over 1,000 annotated small molecules for their ability to reduce ATZ polymer load. Seven hits from the initial single concentration screen were taken forwards and tested with multiple concentrations to measure dose dependence manner in three additional patient iPS lines. Based on the data from these three rounds of testing and because of its clinical relevance in other related disease areas, LRRK2 was selected for further investigation. We began by finding that the LRRK2^−/−^ x ATZ mouse carries less ATZ polymers and shows less liver damage (fibrosis) compared to controls. Next, we saw that a collection of commercially purchased LRRK2 chemical inhibitors could similarly (though to different degrees of efficiency) cause polymer reduction across both iPS and CHO cell models. One of the most efficacious small molecules tested, CZC-25146, a LRRK2 inhibitor with favourable biodistribution properties for targeting the liver, was selected for follow on studies in the ATZ mouse. There it reduced polymer load and inflammation after only 14 days oral treatment. Finally, we confirmed the mechanism of action of CZC-25146 to be via inhibition of LRRK2 kinase activity and augmentation of autophagy. Cumulatively, these data support the idea that LRRK2 inhibitors activate autophagic clearance of ATZ polymers from hepatocytes and that CZC-25146 along with related LRRK2 inhibitors may represent a new therapeutic approach for the liver disease associated with α-1-antitrypsin (z).

Liver disease represents an increasing global health challenge. ATZ associated liver disease is caused by a gain-of-function toxicity arising from AAT polymer accumulation. Recent developments to suppress AAT protein production via RNAi *(29)* or replace missing AAT via donor plasma infusions *(30)* are promising and have progressed to clinical trials but are unlikely to represent a complete longer term solution for patients since these approaches cannot address both liver and lung pathologies concomitantly. Instead, a therapy that is able to reduce ATZ polymers and restore the balance of normal AAT secretion from the liver is most needed. Gene editing to correct underlying mutations in the *SERPINA1* gene or cell therapy could solve this problem but there are currently no technologies advanced enough to facilitate sufficient levels of genetic correction or cellular engraftment needed in order to achieve clinically relevant outcomes. In the interim, a small molecule approach to either block polymer formation *(31)* or increase polymer degradation represent two of the most promising, potentially complimentary, alternative solutions. Targeting intracellular degradation is particularly appealing since the clinical variations in liver disease severity observed in patients harbouring the same genetic mutation (ranging from no/minimal disease to liver failure requiring organ replacement) may result from variable capacity in the patient’s hepatocytes to degrade misfolded proteins. Although the autophagy pathway has been reported to play an important role in ATZ polymer degradation, the identity of the specific molecular players involved has remained elusive. This may in part explain why attempts to non-specifically augment autophagic flux have so far yielded unsuccessful clinical results (ClinicalTrials.gov identifier NCT01379469) *(32, 33)*.Identifying a hepatocyte specific target to accelerate polymer destruction and restore / maintain normal AAT production may solve this issue but until recently has been hampered by a lack of suitable research tools. Using iPSC technology to engineer hepatocytes from patients with clinically documented ATZ polymer associated liver disease, allowed us for the first time as far as we know, to robustly identify molecular targets for polymer degradation by performing a small molecule screen in a cell model wherein both the genetic defect (Glu-Val 342) and patient specific protein degradation machinery are conserved. Previous disease models which employ overexpression of the mutated human gene in a non-human background (eg mouse, hamster or worm) *(14, 34)* cannot achieve these insights because they are limited by species differences in the host cell machinery. We therefore propose that our approach is the most robust currently available tool for identifying protein degradation targets relevant for patients. Indeed all 7 of our validated hits represent highly interesting molecular targets to follow up on. We chose, however to focus in on LRRK2, a seven domain, 285kDa cytosolic protein implicated in lysosomal degradation *(35, 36)*. Previous work by us and others suggests that patients harboring AAT-Z mutations develop liver disease when their polymer disposal mechanisms (autophagy - lysosomal - degradation pathway; ALP) are overrun. LRRK2 may therefore be a critical but hitherto unrecognized regulator of hepatocyte ALP and that by inhibiting LRRK2 kinase activity we can drive ATZ polymer disposal and ameliorate the associated liver disease. Interestingly, genetic variation at the *LRRK2* locus has been identified as a key contributor to the risk of another protein misfolding disorder, Parkinson’s disease *(36)*. This association led to the development of LRRK2 kinase inhibitors and antisense oligonucleotides as potential new medicines for PD, with leading candidates expected to enter late-stage clinical trials this year *(17)*. Their mechanism of effect however remains incompletely understood. Several studies suggest LRRK2 acts upon different stages of the ALP, with conflicting results on how this is mediated and the net physiological direction. For example, in neurons derived from patients with PD, it was shown that Rab10 is a mediator of LRRK2 kinase activity that negatively regulates lysosomal activity and clearance of alpha-synuclein *(37)* whilst in infected macrophages, LRRK2 regulates the ALP, this time via Rab8A interaction with CHMP4B *(38)*. In contrast, LRRK2 overexpression in kidney and neuroendocrine cells was found to induce autophagy through a Ca^2+^/CaMKK/AMPK pathway *(39)*. These studies support a paradigm whereby LRRK2’s role in ALP is both cell- and disease-specific. Our data suggests both the level of LRRK2 protein and its kinase activity are important regulators of ATZ polymer handling via the ALP in hepatocytes and that targeting LRRK2 could lead to a therapeutic reduction in polymer load with downstream reversal of liver disease. We do not as yet know however what the specific molecular players interacting with LRRK2 to regulate those observations are. Future studies to increase our understanding of how LRRK2 works in this context could therefore help drive new treatment approaches for ATZ as well as other liver diseases such as NAFLD in which protein aggregates and ER stress are postulated to represent fundamental components of a multi-factorial pathology *(6, 40–43)*.

In the short-term, our data support further investigation of the LRRK2 inhibitor, CZC-25146, in ATZ. This compound was previously developed as a therapeutic for Parkinson’s disease but did not progress further because it was found to cross the blood-brain barrier poorly *(19)*. That bio-distribution profile in contrast seems advantageous when seeking to target pathology restricted to the liver and therefore provides further impetus to evaluate this molecule in the context of ATZ therapy. More broadly, we propose LRRK2 inhibitors, especially those already shown to be safe in clinical trials of Parkinson’s disease *(24)*, should be considered to not only help further our mechanistic understanding of ATZ biology in the laboratory but also considered for therapeutic use in the clinic.

## MATERIALS AND METHODS

### Study design

The primary objective of this study was to identify and test new therapeutic targets for alpha-1 antitrypsin deficiency disease. In the first part of the study (Fig. 1), we used a high-throughput annotated small molecule screen with automated image analysis to identify molecular targets capable of reducing ATZ polymer levels in patient derived iPSC-Hepatocytes. We then went on to test our lead molecular hit, LRRK2 in three disease models: genetically modified CHO cells inducibly expressing human polymeric ATZ, patient iPSC-ATZ hepatocytes and genetically modified mice over expressing human polymeric ATZ (Fig. 2). In the second part of the study (Fig. 3 and 4), we selected one LRRK2 inhibitor, CZC-25146, for further validation and demonstrated its therapeutic potential using the iPSC (Fig. 3) and mouse models (Fig. 4). Finally, in the last part of the study (Fig. 5) we investigated the mechanism of action with a specific focus on the autophagy degradation pathway.

The design and modality of the mouse treatment experiments are described below. PiZ mouse model was used in the proof-of-concept study, it is a transgenic model with the PiZ variant of the human A1AT gene introduced into the germline of mice to exhibit the ATZ accumulation within the ER of hepatocytes and subsequent resultant in liver necrosis and inflammation as in human patient^2^. This model offers excellent A1ATD liver disease phenotypes. The sample sizes were empirically estimated by considering the variations of the results and the statistical power needed while minimizing the number of animals. Mice were randomized to experimental groups at 6 weeks for PASD globule characterization (polymeric A1AT), or 25 weeks for fibrotic study (Picrosirius red staining). End points for experiments with mice were selected in accordance with institutional-approved criteria; fixed time points of analysis shown in the figures indicate time elapsed from the start of the treatment. Studies were not blinded. The number of technical replicates is described in the figure legends and varies among experiments.

### Cells lines

<u>hiPSCs</u> were cultured in Essential 8 medium (Gibco), on Vitronectin (Sigma Aldrich) coated plates, and passaged using ReLeSR (STEMCELL Technologies) every 3-4 days. hiPSCs were differentiated into hiPSC-derived hepatocytes (iHeps) using our previously established protocol *(8)*. In brief, iHeps were differentiated until day 30 prior to use in all experiments, with plating into assay format taking place at day 21.

<u>Tetracycline-inducible (Tet-ON) CHO-K1</u> cell expressing polymeric A1AT E324K (gift from Lomas Lab) were maintained as previously described *(44)*. CHO-K1 cells were co-treated using 1μg/mL doxycycline (Takara Bio Europe, Clontech Inc.) along with the small molecules for experiments as described in main text.

<u>LC3-GFP fibroblasts</u> were gifted from the Carlton Lab and were analyzed using both the Operetta CLS for quantitative assessment, and the SP8 confocal (Leica) for qualitative assessment. Fibroblasts were cultured in DMEM (Gibco) +10% FBS (Sigma Aldrich) + 1% Penicillin & Streptomycin (Sigma Aldrich) and passaged once a week using Trypsin (sigma). For experimentation, fibroblasts were cultured in OptiMEM (ThermoFisher, 31985062) +/- test compounds, without any FBS. Data analysis was conducted on Harmony.

### Annotated Compound Screen

The annotated compound screen was conducted by the National Phenotypic Screening Center, using a compound library of known molecules with annotated targets, provided by Janssen Research and Development. A day before cell seeding, 384-well plates were coated with laminin-411 (LN411-03, Biolamina) at 3.75μg/cm^2^. The A1ATD and healthy control iPSC-hepatocytes used in the screen were commercially sourced (Definigen), thawed in Def-HEP thawing media (Definigen), and plated at a density of 20,000 cells per well. Subsequently, the cells were maintained in Def-HEP Recovery & Maintenance Medium (Definigen) until day 13 post-seeding. JANUS G3 (Perkin Elmer) automated liquid handling system was used for media changes every two days until day 13, then for test compound dispensing. A library of 1,041 annotated small molecules were all dissolved in DMSO and tested at a final concentration of 10μM for each compound. The test compounds were left for 24 hours, with each compound being tested in duplicate in separate plates. Subsequently, the wells were washed, fixed with 4% paraformaldehyde, and permeabilized with 0.5% Triton X for 5 minutes. The samples were then blocked in 3% donkey serum in 1% BSA for 1 hour and incubated with primary antibodies at 4*C overnight. Polymeric alpha-1 antitrypsin (A1AT) was assessed using the 2C1 monoclonal antibody (1:250 dilution; gifted by the Lomas lab) and Alexa-488 donkey anti-mouse secondary antibody (1:500 dilution; A21202, Life Technologies). Image acquisition was performed on an In Cell 6000 Analyzer (Cytiva), and image analysis performed with Columbus software (Perkin Elmer). The 2C1 signal was quantified by detection of the nuclei (DAPI^+^ objects), then generating a ring-like structure by shape dilation by 2.5μm. The signal intensity was measured in the ring region to obtain the average polymer load measurement per cell in the well. An average of all cells per well was used in the downstream analyses – fold-difference to the untreated A1ATD cells and z-score calculation for each well. The expression of albumin was assessed for quality control purposes and was measured using the anti-albumin goat monoclonal antibody (1:500 dilution; A80-129A, Bethyl) and Alexa-647 donkey anti-goat secondary antibody (1:500 dilution; A21447, LifeTechnologies). Compounds that led to cell death, determined by a dramatic reduction in DAPI^+^ objects, or loss of albumin expression, were removed and excluded from the analysis. Hits were defined as compounds with a 2C1 z-score in the vicinity or lower than 1.92 (95^th^ percentile of healthy control cells) were selected for a confirmatory screen. Fourteen hits selected in the primary screen were advanced onto confirmatory, dose-dependent studies. The compounds were tested in triplicates at two-fold serial dilution doses ranging from 20μM to 156nM. This secondary screen was performed on 384-well plates using the immunofluorescence methodology used in the primary screen. The 2C1 readouts were normalized to the DMSO-treated A1ATD cells and the seven compounds showing dose response and no severe cytoxicity (<75%) were selected for further validation.

### Validation of Compound Screen

CZC-25146 (Tocris, 6071), Flurandrenolide (Sigma, 1284000), Fenofibrate (Sigma, F6020), NU6140 (Sigma, SML1323), 4-Chloro-2-Methylthiopyrimidine (Sigma, 145289), GW3965 (Sigma, G6295), 1-9-Pyrazoloanthrone (Sigma, S5567), CZC-542525 (Sigma, S6534), MLi-2 (Sigma, S9694), GNE-9605 (Sigma, S7368), GNE-0877 (Sigma, S7367) and GNE-7915 (Sigma, S7528) were tested on day 30 ATZ iHeps for either 24 or 48 hours, with polymeric A1AT assessed using the polymer-specific antibody 2c1 (gifted by the Lomas Lab) and viability assessed using the Presto Blue Viability Assay (ThermoFisher), conducted according to manufacturer’s instructions. The appropriate dose response starting point for each compound was determined by using the manufacturer’s solubility data and calculating the top concentration which could be used without exceeding 0.1% DMSO in the culture. Antibodies used include: LRRK2 (Abcam, ab133474), total hA1AT (Abcam, ab9373), human ALB (Bethyl Laboratories, A80-129A). A1AT ELISA (Abcam, ab108798) was used according to the manufacturer’s instructions. High-content imaging was conducted using the Operetta CLS (Perkin Elmer) and data was analyzed using Harmony software (Perkin Elmer). Differential gene expression was detected by quantitative reverse transcription polymerase chain reaction (qRT-PCR). RNA was extracted using RNeasy mini kit (Qiagen, 74104) according to the manufacturer’s instructions. Subsequently, qPCR was conducted using CFX Connect Real-Time PCR Detection System (BioRad). All qPCR data was normalized to GAPDH and RPL13 which are used as housekeepers. Table S1 contains all the primer sequences used during this study.

### Pre-clinical mouse study

Ethical compliance was obtained prior to working with all mouse models. The PiZ mouse was administered CZC-25146 at 250mg/kg dissolved in pure DMSO, administered for 14 days consecutively by oral gavage. Mice were culled after 14 days and each lobe of the liver was separated independently. For IHC characterization, all samples were washed thrice with PBS, followed by a 24-hour fixation with 10% formalin. Samples were then washed again 3x with PBS, before being processed, embedded into paraffin, sliced, and stained. The antibodies/stains used were: PASD (Sigma Aldrich, 395B-1KT), LC3 (Abcam, ab48394), Picrosirius Red (Abcam, ab246832), Ki67 (Abcam, ab15580). *In vivo* image analysis was conducted by taking 3 5x shots of each liver lobe, equally distributed across each lobe (15 images per mouse). The images were analyzed using ImageJ. Western Blot analysis was prepared by homogenizing the frozen liver tissue in Cell Lysis Buffer (Cell Signaling Technology, 9803) supplemented with phosphatase inhibitor (#78427, ThermoFisher) and protease inhibitor (ThermoFisher, 78429) using TissueLyzer II (Qiagen) for 30 min. Protein concentrations were normalized using a Pierce™ BCA protein assay kit (ThermoFisher) according to the manufacturer’s instructions. Processed proteins samples were run on 3–8% NuPAGE Tris-Acetate 1.5mM pre-cast gels (ThermoFisher, EA0378BOX) and primary antibodies used were: anti-phospho-Ser935 LRRK2 [UDD2 10(12)] (Abcam, ab133450) or anti-LRRK2 [MJFF2] (Abcam, ab133474). The secondary antibody was IR Dye 680RD (Licor, 962-68073). Blots were imaged using ChemiDoc MP (Biorad). Quantitation of the bands was performed using Image Studio (Licor).

## Supplementary Materials

**Fig. S1:**
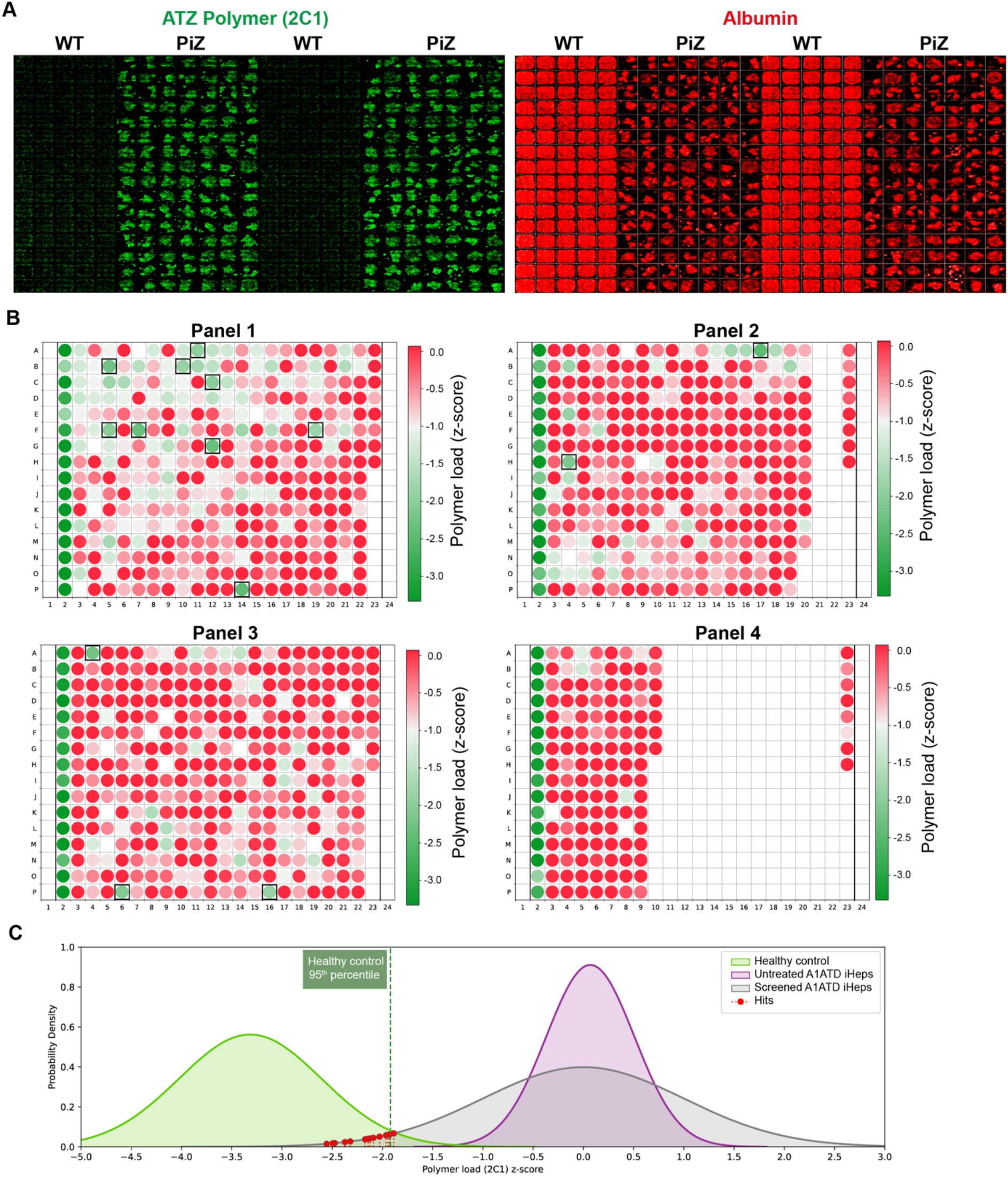
High-throughput screen approach. **(A)** Representative images from WT and PiZ hiPSC-derived hepatocytes in 384 well view showing the 2C1 (disease) and albumin (hepatic) expression. **(B)** Four panels of small molecules (1,041 in total) were used to screen A1ATD plated in a 384 well plate format in duplicates. Each well shown visualizes a mean 2C1 signal of two replicates. Top hits (14) showing the lowest Z-scores were selected for further validation. **(C)** Gaussian curves were plotted to visualize the position of the hits relative to the healthy control (n=64; green) and untreated A1ATD iHeps (n=32; purple). The hits (red) appear to dramatically shift the 2C1 signal towards the range of the healthy control, thus restoring the healthy AAT phenotype in the A1ATD cells.

**Fig. S2:**
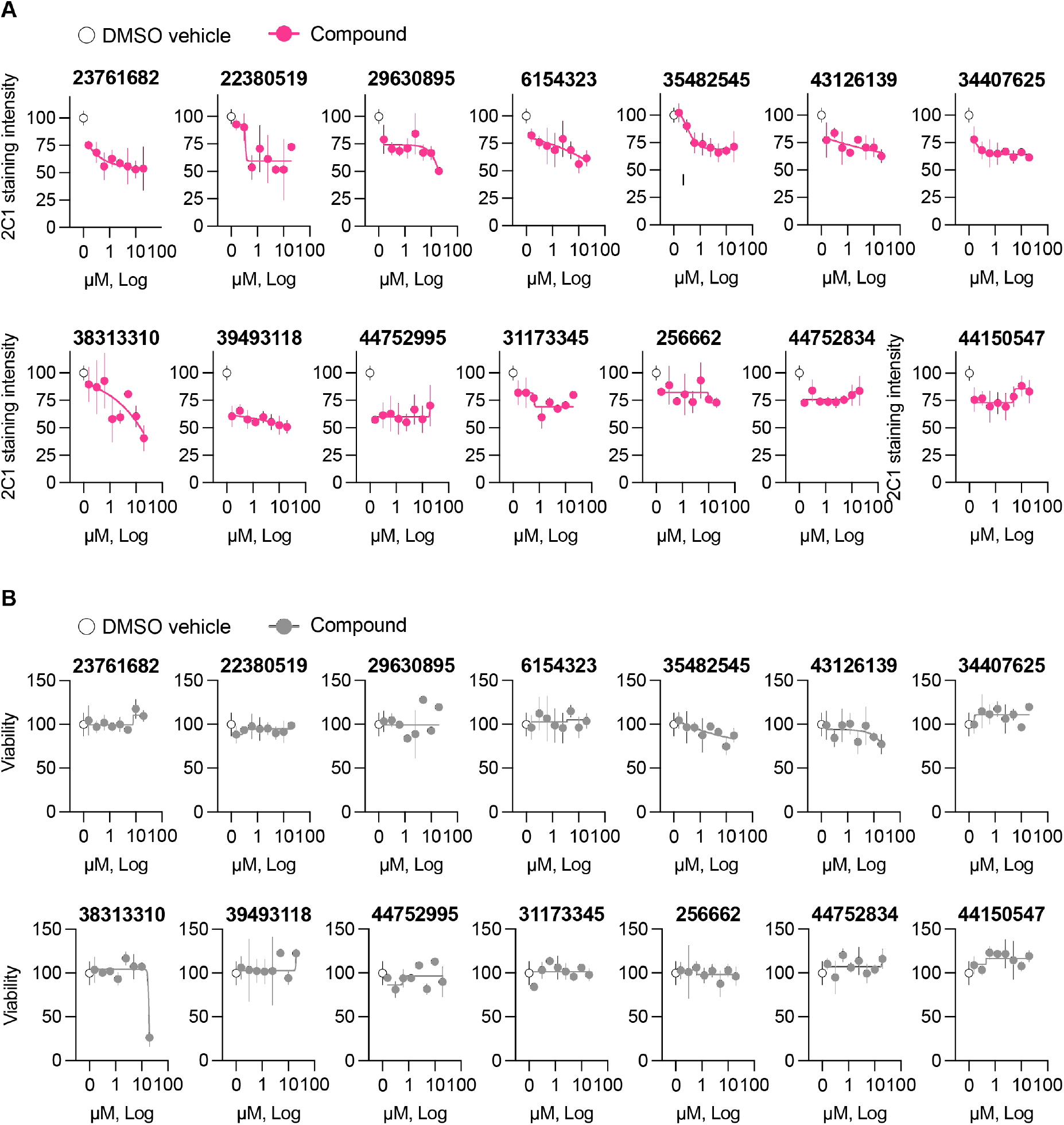
Confirmatory dose-dependent study on the top 14 hits. **(A)** The effectiveness of small molecules is determined by the reduction of 2C1 staining signal (ATZ polymer). (**B**) The safety of small molecules is evaluated by the viability assay by Presto Blue.

**Fig. S3:**
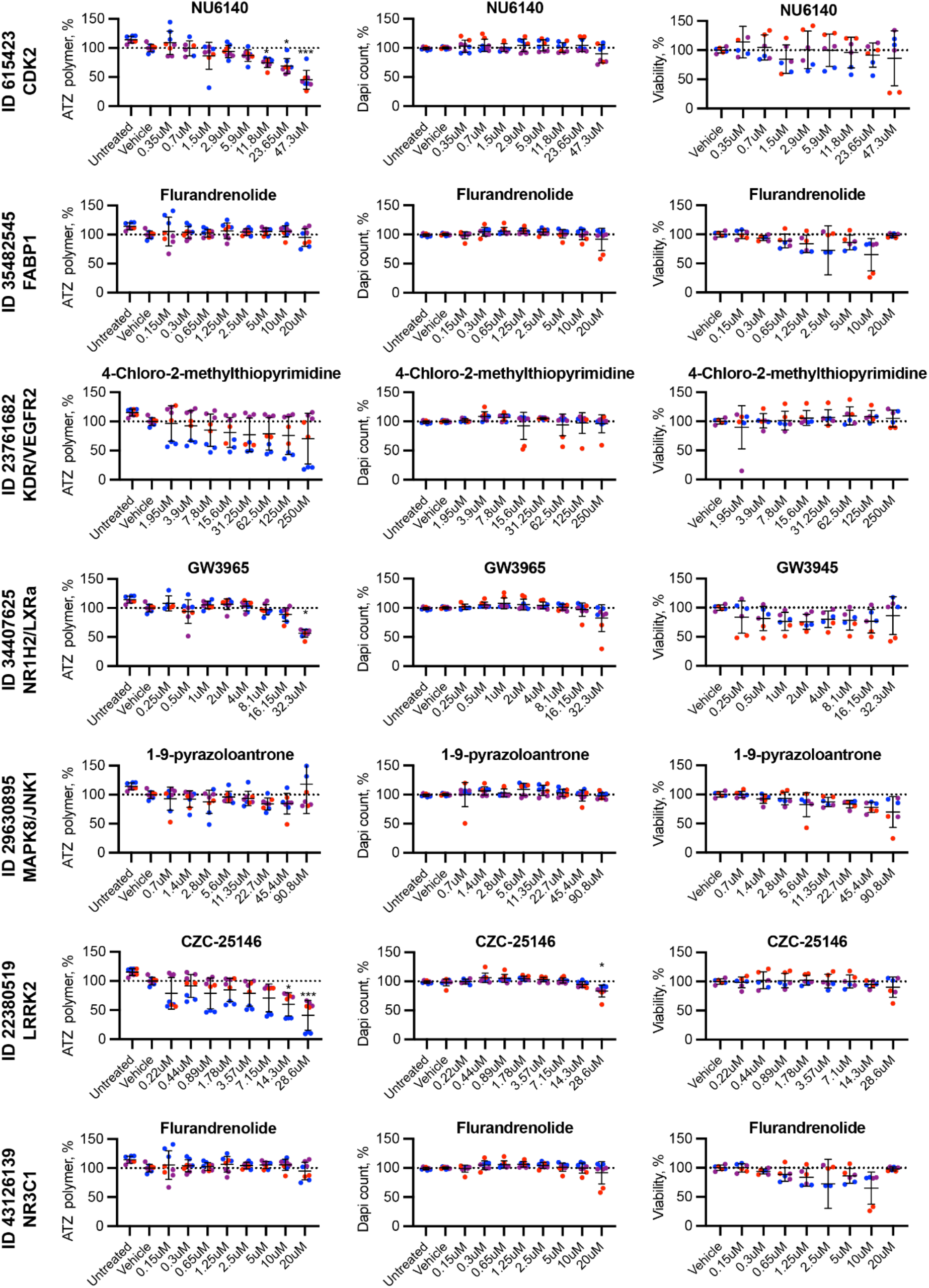
Top 7 putative targets validated using commercially available small molecules. The effectiveness of the small molecules is determined by the reduction of ATZ polymer (right column). The viability of the small molecules is evaluated by the DAPI + object counting (middle column) and the viability assay by Presto Blue (right column). The color of the dot represents each A1ATD patient iPSC line. Each dot represents one batch of hiPSC-hepatocytes. Statistical analysis involved a Kolmogorov-Smirnov test for normality, followed by Kruskal-Wallis test, followed by a Dunn’s multiple comparison test. *p<0.05, **p<0.01, ***p<0.001, ****p<0.0001.

**Fig. S4:**
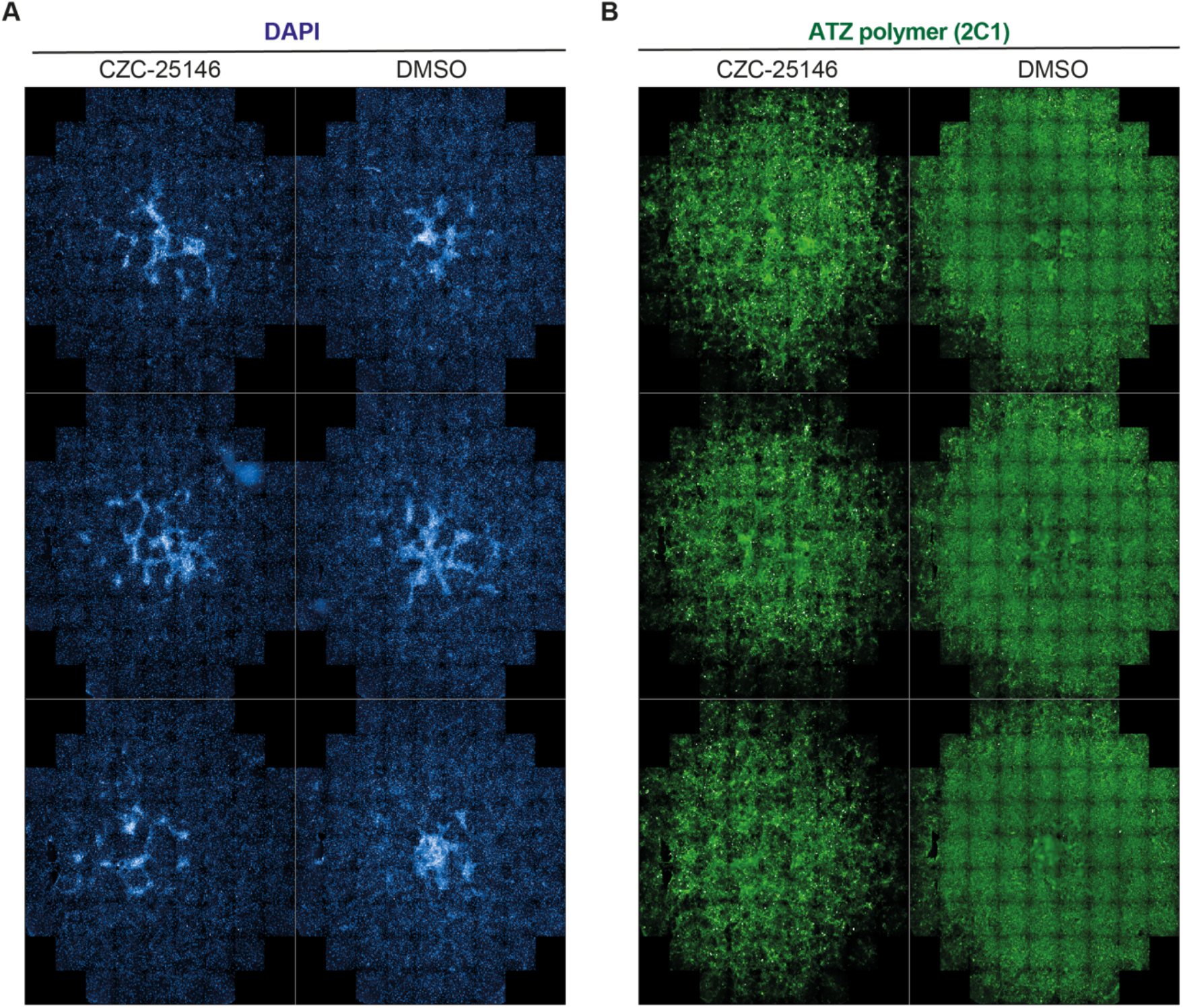
Whole well views of CZC-25146 polymer-reduction phenotype in A1ATD patient-derived hiPSC-hepatocytes. Full 96-well views of CZC and control treatment, after 48 hours of treatment. Each well contains approximately 50,000 PiZ hiPSC-derived hepatocytes. Images taken on Operetta CLS.

**Fig. S5:**
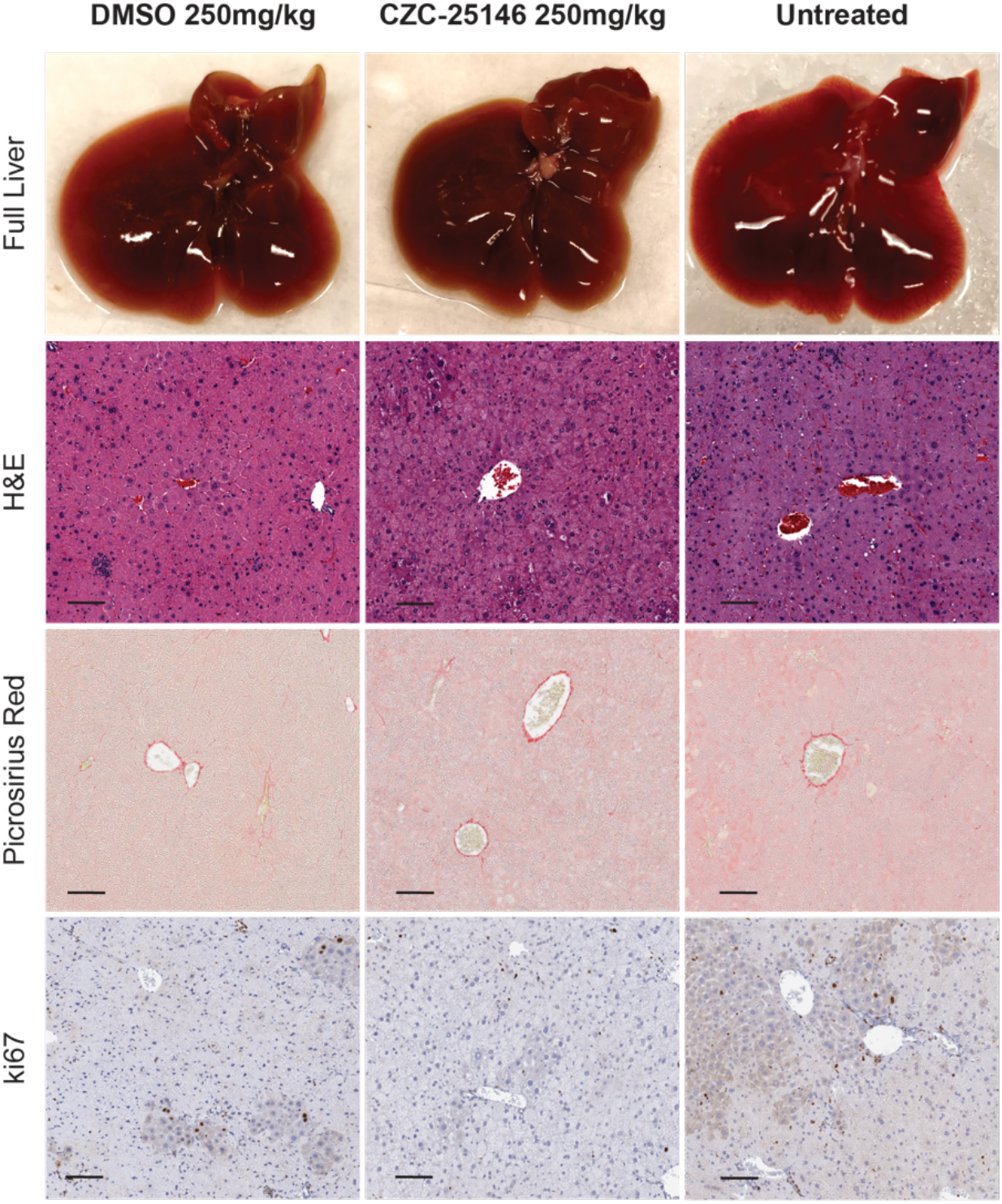
Gross anatomy and IHC staining of the liver treated with CZC-25146 and vehicle control DMSO at 250mg/kg for 14 consecutively days via oral gavage.

**Fig. S6:**
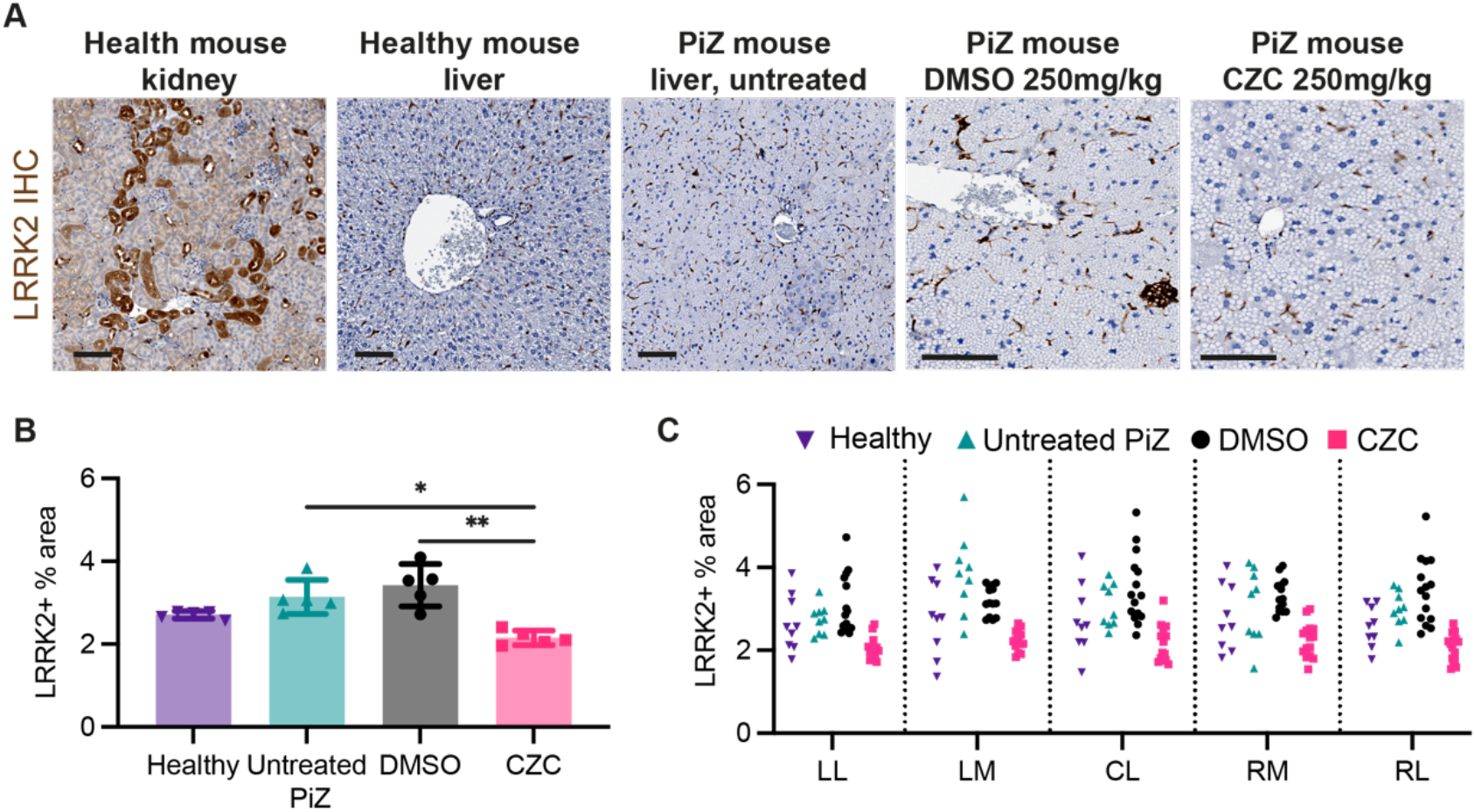
Investigating the expression of LRRK2 *in vivo*. **(A)** Representative images of LRRK2 IHC in healthy mouse kidney used as a positive control for LRRK2 IHC staining, healthy and PiZ mouse livers, PiZ mouse livers treated with DMSO and CZC-25146 at 250mg/kg. All mice used were exactly 6 weeks old and all males. Scale bar, 100μm. LRRK2 IHC quantification in the liver, with all the data pooled (**B**) and the results of each lobe separated (**C**). Quantification was conducted using ImageJ. N= 3 mice per group. Statistical analysis by Ordinary ANOVA test, followed by Dunnett’s multiple comparisons test. *p<0.05, **p<0.01, ***p<0.001, ****p<0.0001.

**Table S1:**
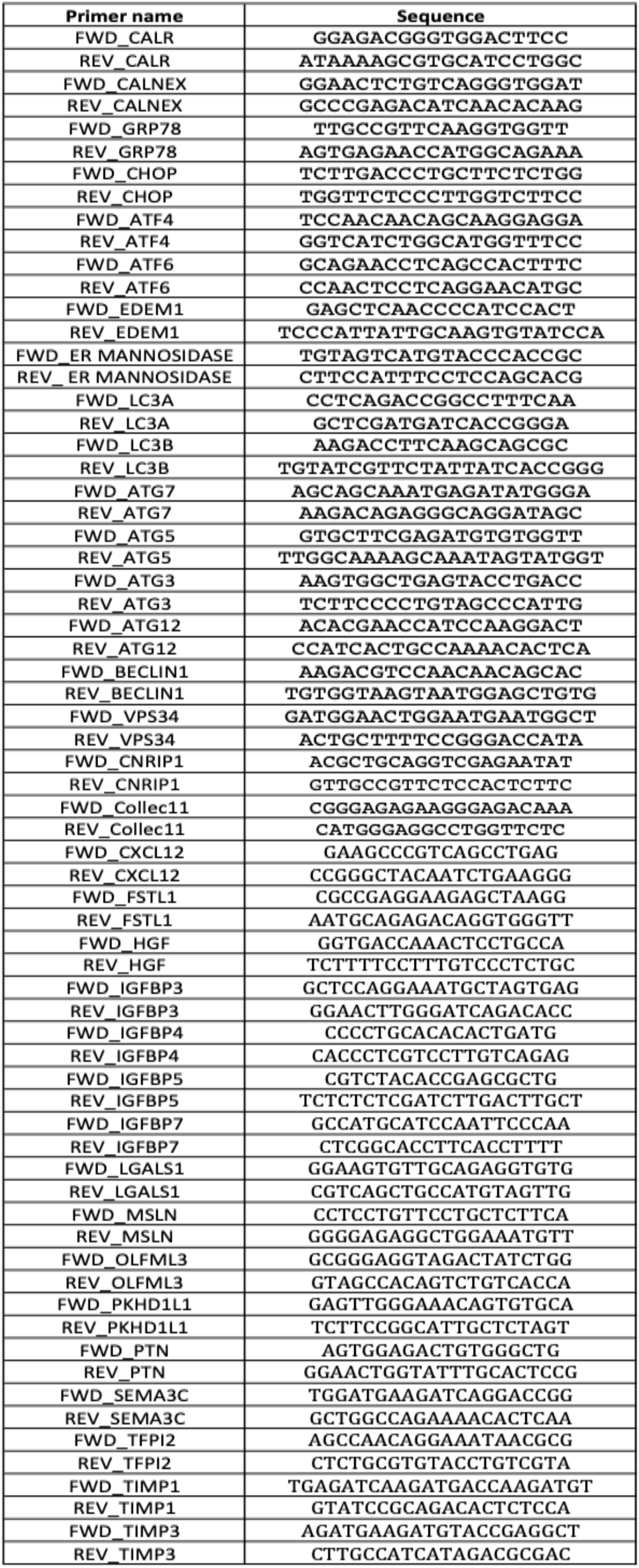
Primer sequences for RT-qPCR assay.

## Acknowledgments

We would like to thank Prof Fiona Watt and Dr. Benedict Oules for reagents and support.

## Author contributions

Conceptualization: STR, CM, DK, SSN

Methodology: STR, SS, DE, DK, SSN

Investigation: STR, CM

Visualization: DK, SSN, AMS,

Funding acquisition: STR, CM

Project administration: DK, SSN

Supervision: STR, CM, DD, DE,

Writing – original draft: SSN, DK

